# Risk preferences depend on environmental richness in rats performing a patch-foraging task

**DOI:** 10.1101/2025.02.04.636458

**Authors:** Marissa Garcia, Andrew M. Wikenheiser

## Abstract

Risk, or variance over outcomes, features prominently in many decisions, but the factors that determine when and how risk modulates decision making remain unclear. We tested how risk affected rats’ strategies for exploiting a diminishing food source as the overall richness of the environment was manipulated. Long-Evans rats earned food by sequentially visiting two foraging patches with different reward schedules—a low-variance standard option and a high-variance risky option—that provided the same average rate of reward. When rats switched between options, they encountered either a long or short delay during which no food was available to simulate the cost of travelling between patches. When the travel delay was short rats allocated more time to the low-variance standard reward schedule than the high-variance risky option. When the travel time was long rats spent the same amount of time in risky and standard patches. Consistent with previous work, rats “overharvested” patches, remaining for longer than the optimal patch residence duration. Overharvesting was prevalent in both risky and standard patches, and the magnitude of overharvesting increased with successive visits to the same patch, suggesting that overharvesting was not driven by uncertainty about the reward statistics of patches.

## Introduction

The consequences of our choices are rarely guaranteed. The same option selected under similar circumstances might nevertheless result in a different outcome. This sort of variance in outcomes—also known as risk—is ubiquitous in nature^1–3^. For example, a foraging animal searching in the same location on successive days might find different amounts of food on each visit. Some sources of variability follow predictable patterns (i.e. the seasonality of fruit-bearing trees), while others may be difficult or impossible to anticipate (i.e. sudden changes in weather or competition from other foragers)^4^.

Risk affects decision making in at least two ways. First, agents may have risk preferences that cause them to favor or avoid risky options^5^. In the laboratory, risk preferences can be measured by examining decisions involving options that are matched in their average return but that differ in their variability. For instance, a non-risky option might offer one unit of food each time it is chosen, while a risky option might yield either two or zero units of food with equal probability. When average returns are equal, a preference for the more variable option is described as risk seeking, a preference for the less variable option is described as risk averse, and indifference between the two is described as risk neutrality. Risk preferences have been assessed in a range of species, and evidence for risk-seeking behavior has frequently been observed^6–11^. However, it is clear that contextual factors can influence risk preferences. For instance, pairing choice outcomes with sound and light cues made rats^12^ and humans^13^ less likely to prefer an advantageous low-variance option to a disadvantageous risky one^14^. Similarly, while most studies have found that Rhesus macaques are risk-seeking on binary choice tasks^8,10,15,16^, testing them on a more naturalistic foraging problem revealed risk-averse attittudes^17^.

A second way that risk affects decision making is by impacting how agents learn from experience. Error-correcting learning mechanisms like the Rescorla-Wagner model^18^ and temporal-difference reinforcement learning^19^ use prediction errors—the discrepancy between the predicted and obtained outcomes on each trial—to update the value of chosen options. During learning prediction errors steadily decrease in magnitude each time a deterministic option is selected.

Risky options are variable, and therefore result in persistently larger prediction errors and slower learning. Thus, for risky and deterministic options with the same average outcome magnitude, uncertainty about the true value of the risky option remains greater for longer during learning.

Uncertainty has been posited to play an especially important role in determining behavior on patch foraging tasks, where subjects must decide how long to remain in one location (or patch) as the rate of reward decreases^1,20,21^. The classic solution to this problem—the marginal value theorem— states that a forager should remain in its current patch until the local rate of reward within the patch falls to the average long-term rate available in the environment under optimal behavior^22,23^. Interestingly, foragers across a range of species tend to “overharvest” patches, remaining longer than the optimal residence times predicted by the marginal value theorem^24–28^. Recent proposals rationalize overharvesting as an adaptive response to uncertainty^28,29^. These models posit that foraging algorithms simultaneously prioritize earning reward and reducing uncertainty about patch quality. Under this assumption residence times longer than those predicted by the marginal value theorem may become optimal, because longer visits provide more opportunities to learn an accurate representation of the patch’s reward statistics. This framework has implications for risky foraging scenarios: given two patches matched in average rate of reward but differing in risk, the variability in returns from the risky patch should impede the forager’s ability to learn the underlying reward statistics. Thus, if uncertainty is one cause of overstaying, a risky patch should—other things equal—induce longer residence times than a less variable patch.

Here, we used a spatial patch-leaving task^30^ to investigate how risk influences foraging decisions in rats. By matching the average reward rate of risky and non-risky patches, we were able to assess risk preferences by comparing how long rats spent in each type of patch. We also tested the hypothesis that uncertainty influences patch residence time by examining patch-leaving decisions on a trial-by-trial basis in risky and non-risky patches.

## Results

### The risky-foraging task

Rats performed a previously-described behavioral task^30^ in which they decided how long to exploit a diminishing food resource. Rats could earn sucrose pellets in two open-field arenas (“patches”) which were connected to one another by a corridor. Motorized doors at each end of the corridor controlled rats’ access to the foraging patches (fig. 1a). While rats occupied a patch, food pellets were dispensed from above following a schedule in which reward rate decreased over time. Rats could exit the patch by retreating to the corridor; after leaving a patch, rats were confined to the corridor for a delay during which no food was available. This “travel time” delay was either long (30 s) or short (5 s), and was fixed for the duration of the session. After the delay elapsed, the door to the next patch opened, and rats were free to enter and forage as before. This sequence repeated for the duration of behavioral sessions.

**Figure 1.**
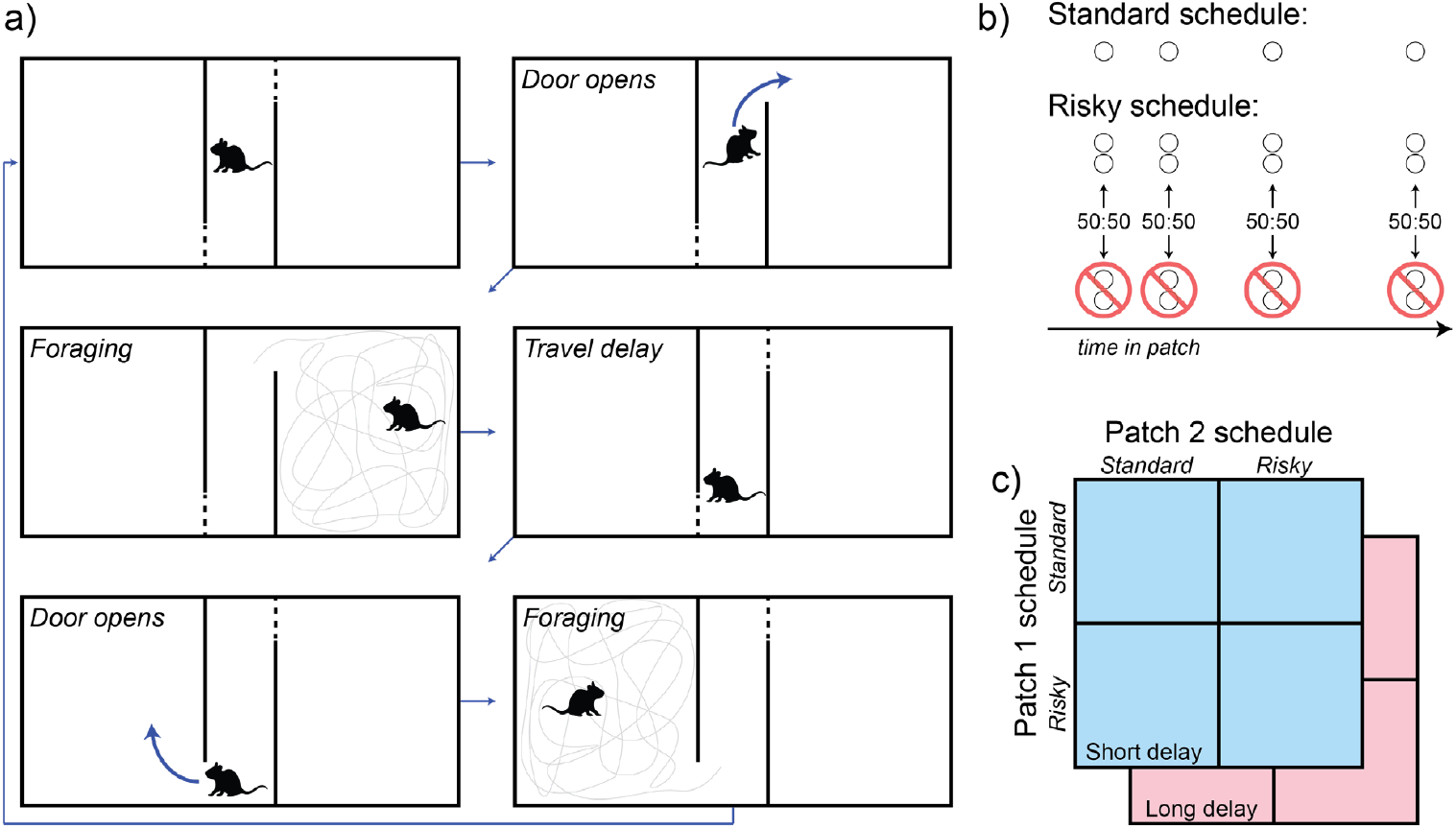
The risky foraging task. (a) Each behavioral session began with the rat in the travel corridor. Next, a randomly-selected door opened, allowing the rat to access one of two foraging patches. The rat chose how long to remain in the foraging patch, collecting pellets that fell into the enclosure from above, according to the reward schedule assigned to that patch. Re-entering the travel corridor caused the door to close behind the rat and remain closed for the travel time duration, after which the door to the opposite patch opened. (b) A reward schedule was assigned to each patch for the duration of each session. In the standard schedule, reward rate decreased exponentially over time while the rat remained in the patch. The risky reward schedule was created by beginning with the standard schedule and randomly omitting half of scheduled pellet deliveries, but doubling the number of pellets per delivery. In this way, the average rate of reward was matched across reward schedules, but the variance of the risky schedule was twice that of the standard schedule. Both schedules reset to their maximal reward rate after each travel delay. (c) In each session, one schedule was assigned to each of the patches comprising the apparatus, and a travel time (5 s or 30 s) was selected. Rats were tested on all possible combinations of session parameters in a randomized order.

This task implements the patch-leaving problem described by behavioral ecologists^1,20–22^, which has been studied extensively in psychology and neuroscience^25,31–38^.

We tested rats on two reward schedules (standard and risky; fig. 1b) with matched time-varying reward rates but different levels of variability. In the standard schedule, reward rate decreased exponentially over time, and inter-reward intervals were sampled from a normal distribution with a mean equal to the reciprocal of the patch’s current rate of reward. Each reward delivery consisted of one sucrose pellet. The risky reward schedule was derived by making two modifications to the standard reward schedule: the number of sucrose pellets per reward delivery was doubled, and the probability of reward delivery was halved^17^. Thus, on the risky schedule the increase in reward magnitude was offset by the decrease in reward frequency, such that the average rate of reward was identical to the standard schedule, but the variability was doubled.

Both reward schedules reset to their maximum value when rats re-entered a patch after the travel delay elapsed. Over the course of the experiment, rats experienced all combinations of reward schedules assigned to the two physical patch locations at both levels of switching cost in a random order (fig. 1c).

### Risk preferences depended on the travel time

As intended, the number of pellets rats harvested was more variable on visits to risky patches compared to patches with the standard reward schedule (fig. 2). While the average number of pellets rats earned during visits of different durations was well matched between the standard (fig. 2a, left) and risky (fig. 2a, right) schedules, the variability was greater for the risky schedule. Fig. 2b displays the same data, but with average earnings for visits of different duration plotted against one another for the standard and risky schedules. The points fall close to unity, indicating that the mean rate of earnings was similar across reward schedules, but the vertical error bars (which denote the standard deviation of risky-schedule earnings) are larger for all visit durations than the horizontal error bars (which denote the standard deviation of standard-schedule earnings). These data show that our risk manipulation effectively controlled the variability of earnings on the task without affecting the rate of reward.

**Figure 2.**
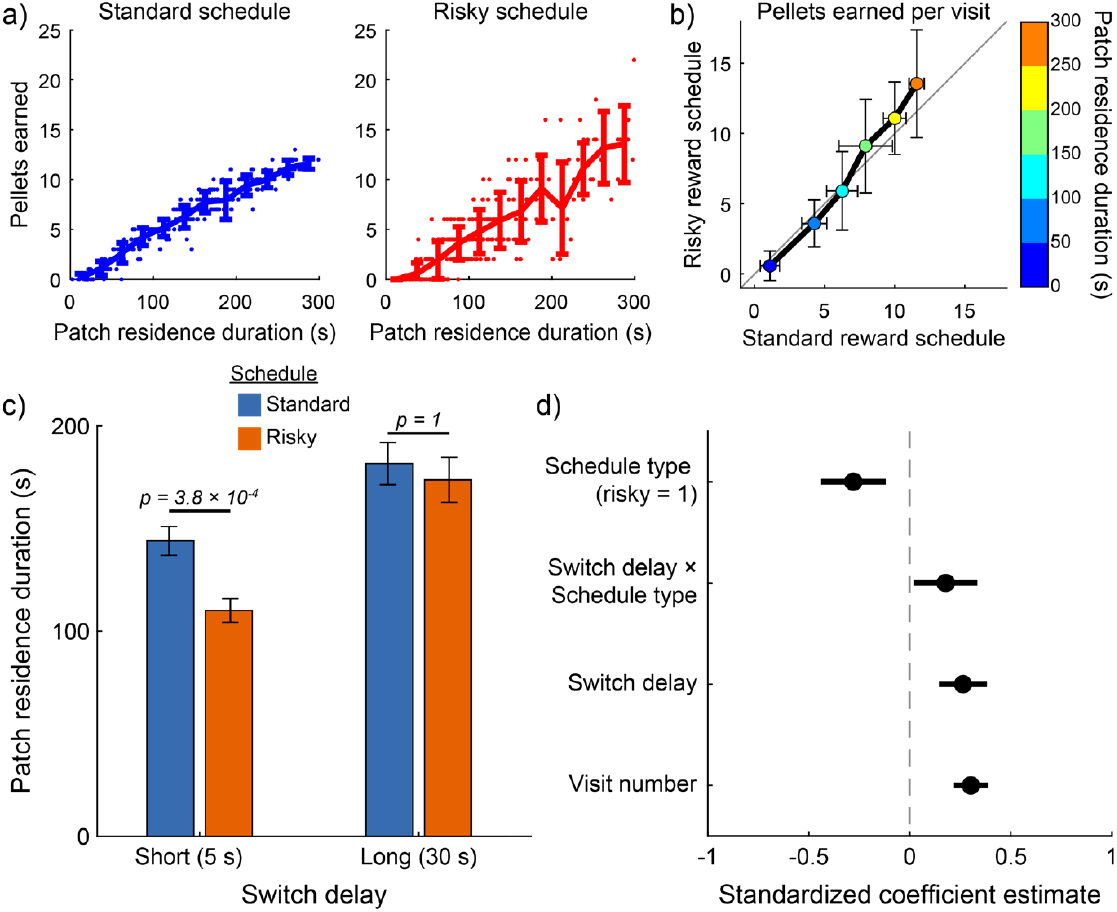
Risk preferences depended on travel time. (a) The number of pellets earned on each visit to a patch is plotted as a function of patch residence duration for standard (left) and risky (right) reward schedules. The line plot shows the average earnings, and error bars depict the standard error of the mean. As intended, rats earned the same number of pellets on average for visits of a given duration, but the variability from visit to visit was greater under the risky reward schedule. (b) Same data, but with visits separated into six duration bins and the mean and standard deviation computed within each bin, separately for standard and risky reward schedules. Points fall near unity (black line), indicating that average earnings were similar between the schedules, but the variability of the random schedule (vertical error bars) exceeds the variability of the standard reward schedule (horizontal error bars) at every duration. (c) Patch residence durations were greater when the travel time was long compared to short travel time sessions. Rats spent less time foraging in patches assigned a risky reward schedule when the travel time was short, but were indifferent between reward schedules when the travel time was long. (d) Coefficient estimates (± 95% confidence interval) are plotted for the mixed-effects model analyzing patch residence duration.

Consistent with models of optimal foraging, rats remained in patches longer when the travel time was long (fig. 2c). However, travel time and reward schedule interacted (fig. 2d; β_travel time * reward schedule_ = 0.37; P = 0.01), suggesting that the effect of risk on residence time differed depending on the travel time. We conducted post-hoc tests to examine how risk impacted residence time at long and short travel times. Rats were risk averse when the travel time was short (P = 0.03, t_209_ = 2.60; paired t-test with Holm-Bonferroni correction for multiple comparisons), spending less time foraging in risky patches compared to patches with the standard reward schedule. Rats were indifferent to risk when the travel time was long (P = 0.81, t_128_ = 0.24; paired t-test with Holm-Bonferroni correction for multiple comparisons). The long travel time significantly increased residence duration under both standard (P = 0.04 , t_172_ = 2.33; paired t-test with Holm-Bonferroni correction for multiple comparisons) and risky (P = 3.11×10^-5^, t_165_ = 4.62; paired t-test with Holm-Bonferroni correction for multiple comparisons) reward schedules. The travel time effectively determined the overall richness of the task, significantly influencing rats’ food earnings (pellets earned [mean ± standard deviation] in low switch cost sessions = 70.0 ± 8.6; pellets earned in high switch cost sessions = 61.2 ± 9.3; P = 0.002; t_62_ =3.2; two-sample t-test). Thus, these results show that risk preferences were modulated by environmental richness, with rats showing a greater tolerance for risk when the cost of switching patches was high and overall richness was low.

### Rats over-harvested patches

Visual inspection of the behavioral data showed that rats generally made brief visits to patches early in the session and gradually extended the duration of their visits as the session progressed (fig. 3). Indeed, visit number significantly modulated patch residence duration (β_visit number_ = 0.09; P = 1.30×10^-5^; fig. 2d). This systematic change in visit length might reflect an adaptive process by which rats gradually adjusted their behavior toward the optimal patch residence time. Numerical simulations determined that the optimal patch residence time was 89 s for short travel time sessions, and 147 s when the travel time was long. Rats’ average visit durations exceeded the optimal residence times (fig. 2c). We computed overstaying—the difference between observed and optimal residence durations—stratified by reward schedule and travel time (fig. 4). For all conditions, the 95% confidence interval around average overstaying did not include zero, indicating that rats significantly overstayed patches in all task conditions (fig. 4a). We next examined overstaying as a function of patch visit number (fig. 4b). The systematic increase in residence time that occurred within sessions led to progressively greater overstaying as sessions progressed. Initial visits to patches were near or below the optimal residence duration, while successive visits eventually exceeded the optimal duration, in some conditions nearly doubling it. Importantly, this systematic change in residence time over successive visits was similar across patches with risky and standard reward schedule and at both levels of travel time. In all four conditions, residence duration and visit number were positively correlated (short travel, standard schedule, R = 0.25, P = 7.00×10^-4^; long travel, standard schedule, R = 0.34, P = 5.00×10^-4^; short travel, risky schedule, R = 0.19, P = 6.20×10^-3^; long travel, risky schedule, R = 0.26, P = 0.01).

**Figure 3.**
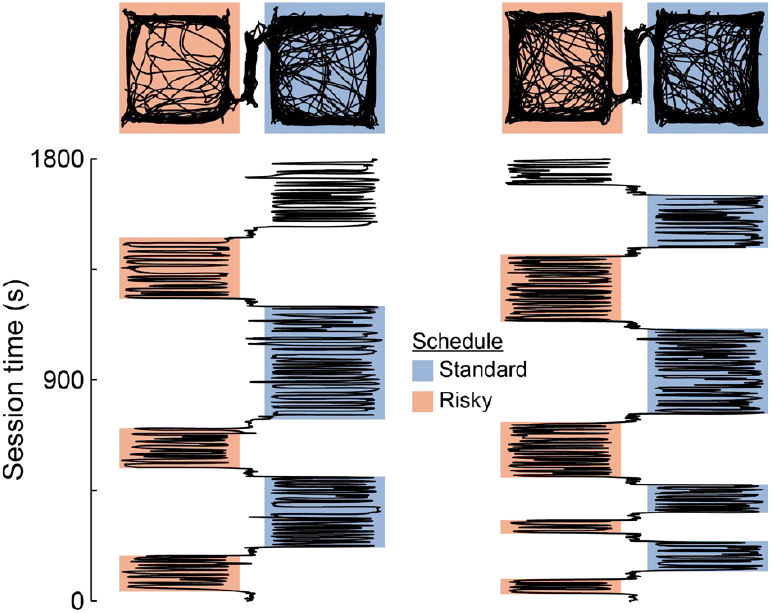
Patch visits increased in duration within sessions. Position-tracking data for two sessions depicts a top-down view of rats’ movements across the apparatus (top), and x-position as a function of time (bottom). The duration of visits to each patch are denoted by shaded blocks, with color indicating the type of reward schedule assigned to the patch. In both sessions, early visits to patches were relatively short, with duration generally increasing for subsequent visits.

**Figure 4.**
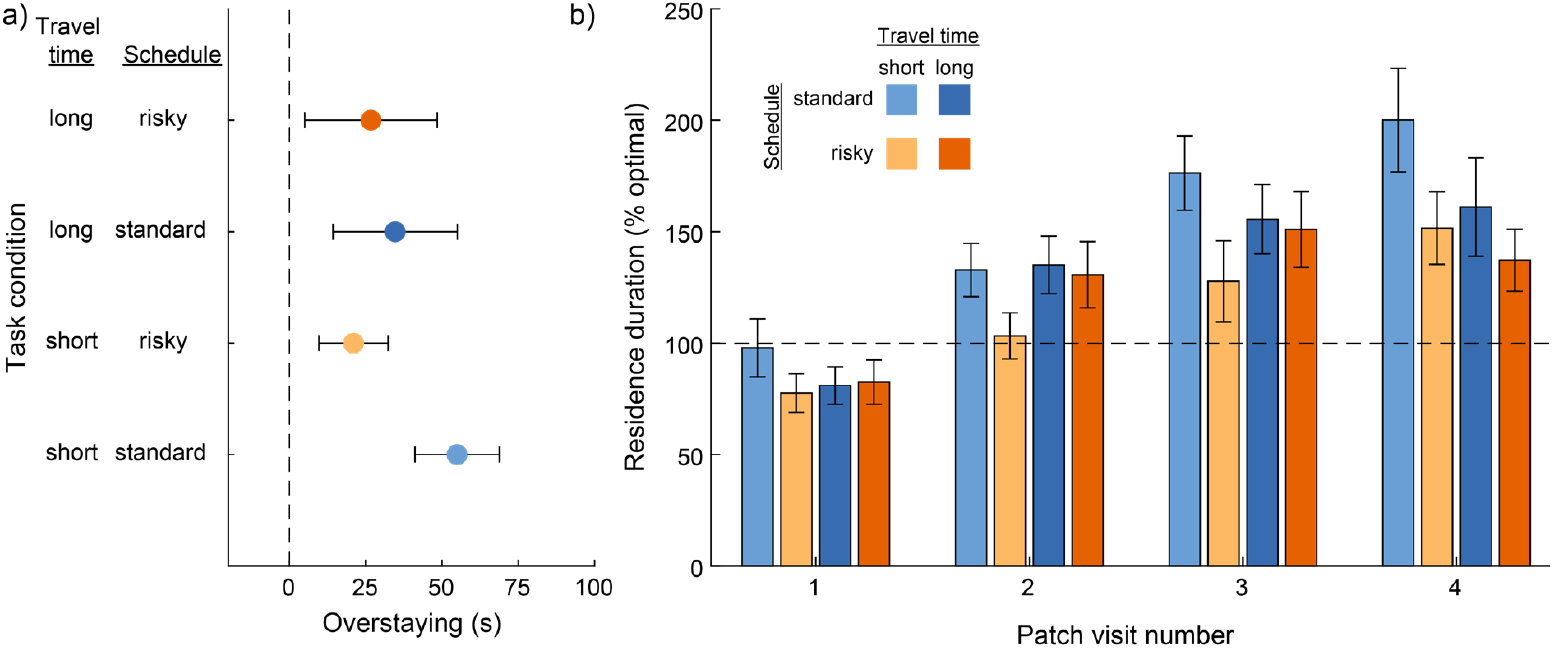
Rats overharvested patches. (a) For all combinations of travel time and reward schedule, overstaying (defined as the difference between the observed and the optimal residence time) was positive, indicating that rats remained in patches longer than the optimal residence time. Error bars depict 95% confidence interval on overstaying for each condition. (b) Residence duration (stratified by travel time and reward schedule, and normalized to the optimal residence duration) is plotted for the first four visits to each patch. For three of four task conditions, the average residence duration on the first patch visit of the session was shorter than the optimal value. Overstaying increased with each subsequent visit for all task conditions, with rats remaining around 50% longer than the optimal residence time by their fourth visit to a patch.

## Discussion

Rats spent less time foraging in risky patches compared to standard patches when the travel time was short, despite the risky and standard reward schedules offering the same average rate of reward. While models of foraging like the marginal value theorem maximize the long-term average rate of reward without considering the higher moments of reward rate^1,22^, risk-sensitive foraging models^6,39^ consider both average rate of reward and variability in returns. Our results show that rats are sensitive to variability in outcomes, at least under some testing conditions.

Although rats were risk averse in sessions with a short travel time, they spent equivalent amounts of time in risky and standard patches when the travel time was long. This is consistent with previous work showing that risk preferences can depend on the broader decision making context. A long-standing framework—the energy budget rule—offers a normative account that suggests foragers should be tolerant of risk in a lean environment^40,41^. Under dire circumstances, where starvation is a real possibility, an especially large return from a risky option might provide enough calories for the forager to survive another day, while a smaller but more reliable payout might not^3,6,39,42^. Although empirical support for this idea is mixed, it has motivated a rich body of experimental and theoretical work^6,39^.

Travel time was the major determinant of how environmental richness, or how much food rats could earn on our task. By increasing the cost of switching patches, the long travel time forced rats to remain in patches for a longer duration on each visit, and therefore experience lower rates of reward due to patch depletion. Thus, contrary to the predictions of the energy budget model and the results of a similar experiment in Rhesus macaques^17^, rats on our task became less risk averse when the environmental richness was low. Our study has several notable limitations that should be considered when interpreting these results in the light of risk-sensitive foraging models and the energy budget framework in particular. The level of food deprivation rats in this study experienced was mild compared to what they might experience in nature, and we maintained their body weight with supplemental food outside of the task to hold motivation stable across sessions. Further, the experimental subjects here were inbred laboratory rats purchased from a commercial vendor and housed in relative luxury compared to their wild counterparts. To the extent that risk attitudes and foraging strategies are influenced by any of these factors, our experiments might not constitute a strong test of the risk-sensitive foraging framework as originally conceived. This work does, however, contribute to a large body of evidence showing that risk attitudes are modulated by a wide range of internal and external factors across various species^6,8,11,43–46^. It also bears mentioning that the riskiness of a patch could be manipulated in many ways besides the approach that we chose here. In binary choice tasks, previous investigations have manipulated risk in reward amount and delay to reward separately. Our risky reward schedules manipulated variability in reward size and timing simultaneously, which makes it difficult to unambiguously attribute the observed changes in behavior to either of these factors. Future experiments could more directly dissociate these factors.

As in many previous studies of patch leaving, rats remained in patches longer than the optimal residence time. Interestingly, this tendency to overstay steadily increased throughout the session, with the first visit duration tending to be at or below the optimal residence time. It is unclear what drives this systematic change in foraging behavior, but it was consistent across travel times and reward schedules. It is tempting to view the initial, short patch visits rats made as exploratory; by briefly examining each of the patches, rats might have been confirming the absence of predators or other threats in each location while also gaining partial information about each patch’s reward schedule. Changes in patch residence time as a function of experience have been observed in similar patch foraging tasks with mice^47^, Rhesus macaques^29^, and humans as subjects^28,29^. Interestingly, in all of these cases, patch residence time *decreased* over the course of sessions.

It has been argued that overstaying may arise due to information seeking^28,29,48^, and the decreases in residence time that have been observed in other studies as foragers become more familiar with patches is broadly consistent with that idea. The information-seeking account predicts that risky reward schedules, due to their inherent variability prolonging uncertainty about the true rate of reward, should drive longer residence times than less variable schedules. In fact, rats spent less time in risky patches when the travel time was short, and the same amount of time in risky and standard patches when the travel time as long. This finding, coupled with the fact that overstaying increased as rats became more familiar with (and therefore less uncertain about) patches suggests that other factors may give rise to the overstaying rats exhibit on our version of the patch-leaving task^27,30^. Future work might more directly manipulate rats’ uncertainty about patch reward statistics. For instance, changing the reward schedule mid-session would create a scenario where initial learning and subsequent experience are in direct conflict. Examining such unsignaled changes in patch quality would speak to how rats learn patch reward statistics and how they update their understanding in the face of new information.

## Materials and methods

### Subjects

Four Long-Evans rats (2 female) were purchased from Charles River at 10-12 weeks of age. Rats were food restricted to maintain them between 85% and 90% of their free-feeding weight; the food rats earned on the foraging task was insufficient for them to maintain weight in this range, so supplemental rodent chow was provided each day, with the amount titrated based on how much food rats earned during task performance. Rats were housed singly under a standard 12-hour light cycle. We did not track levels of circulating sex hormones in either male or female rats. All procedures followed the NIH Guide for the Care and Use of Laboratory Animals and were approved by the Chancellor’s Animal Research Committee at the University of California, Los Angeles.

### Handling and habituation

Rats were handled for approximately 10 minutes per day to habituate them to contact with the experimenter for 7 days before testing sessions began. During the handling period, rats were also exposed to the foraging apparatus daily for approximately 10 minutes. Both doors were set to the open position and food pellets were randomly scattered across the foraging patches during these habituation sessions. By the end of the handling period rats were observed to explore all portions of the apparatus and consume the food pellets they encountered in the patches, and so were deemed ready for experimental sessions to begin.

### Behavioral task

Detailed methods have been reported previously^30^. Briefly, the testing apparatus comprised of two square-shaped open field arenas (77 cm) connected to one another by a corridor (25 × 77 cm). Doors at each end of the corridor were controlled by stepper motors. An IR-sensitive camera controlled by Neuromotive (Blackrock) tracked rats in real time. Sucrose pellets could be delivered into each patch from 40-mg food pellet dispensers (Med Associates) mounted above the arenas.

Rats were tested once each day in 30-minute behavioral sessions. In each session, rats began in the corridor with both doors close. The session began when a randomly-selected door to a patch opened, allowing rat access to one of the foraging arenas. After the rat entered the arena, food pellets were dispensed following the reward schedule assigned to that patch until the rat chose to exit the patch by re-entering the corridor. On leaving, the patch’s door closed behind the rat, confining it to the corridor for the travel time delay assigned to that session. After the travel time elapse, the door to the other patch opened, and the rat was again free to enter and forage. This process of visiting patches in an alternating sequence continued until the session ended.

Rats were tested on two reward schedules. In the standard schedule, the average rate of reward decreased as rats occupied the patch, beginning with an initial rate of 1 pellet every 15 s and decreasing exponentially with a time constant of 300 s. Inter-pellet intervals consistent with this average reward schedule were drawn from a normal distribution (µ = 1/current reward rate; σ = 4 s). Sampling inter-pellet intervals in this way made pellet deliveries unpredictable on each visit to the patch, but consistent with the programmed reward function on average. The reward rate reset to its initial value each time a rat re-entered a patch. The risky reward schedule was programmed by generating a set of pellet delivery times as for the standard schedule and randomly omitting 50% of the pellet deliveries. To maintain the same average rate of reward despite decreasing the probability of pellet deliveries, the magnitude of each pellet delivery was doubled to two sucrose pellets.

Rats were tested on a randomized session sequence that fully counterbalanced reward schedule type (risky or standard), the physical patch location (left or right arena), and travel time (30 s or 5 s). This produced 8 unique combinations of task parameters, and rats were tested on each combination twice, for a sequence of 16 sessions. One session was excluded from analysis because technical issues with the position tracking software resulting in no data being recorded. One rat was accidentally tested on the same session parameters twice; both sessions were included in the final data set. This resulted in a final dataset consisting of 64 behavioral sessions and 578 patch visits.

### Data analysis

All analyses were conducted in Matlab. Patch residence time was defined as the time between patch entry and patch exit. Patch entry was determined by finding the first position tracking sample within the bounds of the foraging patch after the door to the patch opened. Patch exit time was defined as when the rat entered far enough into the travel corridor to trigger the door to close. If the session ended while a rat was foraging in a patch, the in-progress visit was excluded due to uncertainty over how much longer the rat would have remained in the patch.

We fit a linear mixed-effects model^49^ to assess how task parameters affected patch residence time. The formula of the model was:

Patch residence duration ∼ intercept + travel time + reward schedule + visit number + reward schedule * travel time + (1 | subject ID)

The model was fit using the Matlab function *fitlme*. Continuous predictors and the dependent variable were z-scored to facilitate comparison across regression coefficients. Patch visits that deviated from the mean by more than 2 standard deviations (<3% of total patch visits) were considered outliers and excluded from analysis. Visit number was included as a predictor because visual inspection of example sessions (fig. 3) revealed a trend for patch residence duration to increase over the course of sessions. The interaction of schedule type (standard or risky) and travel time was included in the model to test the hypothesis that environmental richness modulated risk preferences given previous theoretical models^6,39^ and empirical reports^17^ raising this possibility. Travel time and visit number were coded as continuous variables, while reward schedule was coded as a categorical variable (risky schedule = 1). A random intercept for each subject was included to account for the repeated-measures design of the experiment. We initially fit a fuller model including all of the covariates described above, as well as sex and physical patch location as categorical fixed effects; coefficient estimates were not significant for either of these factors so they were removed from the final model to conserve statistical power and improve the coefficient estimates for more reliable predictors of patch residence time.

### Determining the optimal residence time

We used numerical simulations to find the residence time in each task condition that maximized total food earnings on average. We simulated an agent that remained in each patch for a fixed residence duration on each visit. Reward schedules for simulated patch visits were constructed as described above, and any pellets delivered within the agent’s residence time for that patch were harvested. Travel time in the simulations was set to either 30 seconds or 5 seconds to simulate long and short travel time sessions. We tested agents that used residence times ranging from 0 to 300 seconds in one-second intervals. The simulation was repeated 10,000 times for each session condition and agent residence time. Because the risk and standard reward schedules were matched in their average rate of reward, the best residence times determined in this way were largely insensitive to risk, differing by ∼1-2 seconds when the simulation procedure described above was repeated multiple times. For simplicity, we averaged across the best residence times determined for risky and non-risky conditions (separately for long and short travel times), and rounded to nearest whole second. Following this approach, the best residence times were determined to be 89 seconds (short travel time) and 147 seconds (long travel time). These values were used for all analyses of overstaying, where overstaying was defined as the difference between the rat’s observed residence time on a given visit and the optimal residence time.

## Author contributions

MG and AMW designed the experiment. MG collected the data. AMW and MG analyzed the data. AMW wrote the paper, with input and edits from MG.

## Acknowledgements

This work was supported by a Whitehall foundation research grant (AMW) and a BBRF Young Investigator Grant (AMW). MG was supported by the Build-PODER program (NIH RL5GM118975) at California State University, Northridge. The authors thank members of the Wikenheiser lab for helpful discussions about this work.

## Data availability statement

Data and code are available at https://zenodo.org/records/14791441

## References

1. Stephens, D. & Krebs, J. Foraging Theory. (Princeton University Press, Princeton N.J., 1986).

2. Kacelnik, A. Normative and descriptive models of decision making: time discounting and risk sensitivity. In Characterizing Human Psychological Adaptations (eds. Bock, G.R. & Cardew, G.) vol. 208 51–66 (Wiley, Chichester UK, 1997).

3. Kacelnik, A. & Bateson, M. Risky Theories — The Effects of Variance on Foraging Decisions. Amer. Zool. 36, 402–434 (1996).

4. Rudebeck, P. H. & Izquierdo, A. Foraging with the frontal cortex: A cross-species evaluation of reward-guided behavior. Neuropsychopharmacol. 47, 134–146 (2022).

5. Von Neumann, J. & Morgenstern, O. Theory of Games and Economic Behavior. xviii, 625 (Princeton University Press, Princeton, NJ, US, 1944).

6. Houston, A. I. & Rosenström, T. H. A critical review of risk-sensitive foraging. Biological Reviews 99, 478–495 (2024).

7. Roitman, J. D. & Roitman, M. F. Risk-preference differentiates orbitofrontal cortex responses to freely chosen reward outcomes. European Journal of Neuroscience (2010).

8. Hayden, B. Y. & Platt, M. L. Temporal Discounting Predicts Risk Sensitivity in Rhesus Macaques. Current Biology 17, 49–53 (2007).

9. So, N.-Y. & Stuphorn, V. Supplementary Eye Field Encodes Option and Action Value for Saccades With Variable Reward. Journal of Neurophysiology 104, 2634–2653 (2010).

10. Xu, E. R. & Kralik, J. D. Risky business: rhesus monkeys exhibit persistent preferences for risky options. Front. Psychol. 5, (2014).

11. Ludvig, E. A., Madan, C. R., Pisklak, J. M. & Spetch, M. L. Reward context determines risky choice in pigeons and humans. Biol Lett 10, 20140451 (2014).

12. Barrus, M. M. & Winstanley, C. A. Dopamine D3 Receptors Modulate the Ability of Win-Paired Cues to Increase Risky Choice in a Rat Gambling Task. J. Neurosci. 36, 785–794 (2016).

13. Cherkasova, M. V. et al. Win-Concurrent Sensory Cues Can Promote Riskier Choice. J. Neurosci. 38, 10362–10370 (2018).

14. Winstanley, C. A. & Hynes, T. J. Clueless about cues: the impact of reward-paired cues on decision making under uncertainty. Current Opinion in Behavioral Sciences 41, 167–174 (2021).

15. O’Neill, M. & Schultz, W. Coding of reward risk by orbitofrontal neurons is mostly distinct from coding of reward value. Neuron 68, 789–800 (2010).

16. Genest, W., Stauffer, W. R. & Schultz, W. Utility functions predict variance and skewness risk preferences in monkeys. Proceedings of the National Academy of Sciences 113, 8402–8407 (2016).

17. Eisenreich*, B. R., Hayden, B. Y. & Zimmermann, J. Macaques are risk-averse in a freely moving foraging task. Sci Rep 9, 15091 (2019).

18. Rescorla, R. A. & Wagner, A. R. A theory of Pavlovian conditioning: Variations in the effectiveness of reinforcement and nonreinforcement. In Classical Conditioning II: Current Research and Theory (eds. Black, A.H. & Prokesy, W.F.) 64–99 (Appleton Century Crofts, New York, 1972).

19. Sutton, R. S. & Barto, A. G. Reinforcement Learning: An Introduction. (MIT Press, Cambridge MA, 1998).

20. MacArthur, R. H. & Pianka, E. R. On optimal use of a patchy environment. The American Naturalist 100, 603–609 (1966).

21. Stephens, D. W. Decision ecology: foraging and the ecology of animal decision making. Cogn Affect Behav Neurosci 8, 475–484 (2008).

22. Charnov, E. L. Optimal foraging, the marginal value theorum. Theoretical Population Biology 9, 129–136 (1976).

23. Kolling, N. & Akam, T. (Reinforcement?) Learning to forage optimally. Current Opinion in Neurobiology 46, 162–169 (2017).

24. Nonacs, P. State dependent behavior and the Marginal Value Theorem. Behavioral Ecology 12, 71–83 (2001).

25. Constantino, S. M. et al. A Neural Mechanism for the Opportunity Cost of Time. 173443 https://www.biorxiv.org/content/10.1101/173443v1 (2017) doi:10.1101/173443.

26. Cash-Padgett, T. & Hayden, B. Behavioural variability contributes to over-staying in patchy foraging. Biology Letters 16, 20190915 (2020).

27. Kendall, R. K. & Wikenheiser, A. M. Quitting while you’re ahead: Patch foraging and temporal cognition. Behavioral Neuroscience 136, 467–478 (2022).

28. Harhen, N. C. & Bornstein, A. M. Overharvesting in human patch foraging reflects rational structure learning and adaptive planning. Proceedings of the National Academy of Sciences 120, e2216524120 (2023).

29. Barack, D. L., Parodi, F., Ludwig, V. & Platt, M. L. Information gathering explains decision dynamics during human and monkey reward foraging. 2023.10.14.562362 Preprint at 10.1101/2023.10.14.562362 (2023).

30. Garcia, M., Gupta, S. & Wikenheiser, A. M. Sex differences in patch-leaving foraging decisions in rats. Oxford Open Neuroscience 2, kvad011 (2023).

31. Baum, W. M. Studying Foraging In The Psychological Laboratory. In Advances in Psychology (ed. Mellgren, R. L.) vol. 13 253–283 (North-Holland, 1983).

32. Shettleworth, S. J. Animals foraging in the lab: Problems and promises. Journal of Experimental Psychology: Animal Behavior Processes 15, 81–87 (1989).

33. Hayden, B. Y., Pearson, J. M. & Platt, M. L. Neuronal basis of sequential foraging decisions in a patchy environment. Nature Neuroscience 14, 4178–4187 (2011).

34. Kolling, N., Behrens, T. E. J., Mars, R. B. & Rushworth, M. F. S. Neural Mechanisms of Foraging. Science 336, 95–98 (2012).

35. Kane, G. A. et al. Increased locus coeruleus tonic activity causes disengagement from a patch-foraging task. Cogn Affect Behav Neurosci 17, 1073–1083 (2017).

36. Mobbs, D., Trimmer, P. C., Blumstein, D. T. & Dayan, P. Foraging for foundations in decision neuroscience: insights from ethology. Nature Reviews Neuroscience 19, 419 (2018).

37. Hall-McMaster, S., Dayan, P. & Schuck, N. W. Control over patch encounters changes foraging behavior. iScience 24, 103005 (2021).

38. Logue, R. S., Garcia, A. K., Bejarano, B. A., Kendall, R. K. & Wikenheiser, A. M. Rats pursue food and leisure following the same rational principles. 2024.12.08.627420 Preprint at 10.1101/2024.12.08.627420 (2024).

39. Kacelnik, A. & El Mouden, C. Triumphs and trials of the risk paradigm. Animal Behaviour 86, 1117–1129 (2013).

40. Stephens, D. W. The logic of risk-sensitive foraging preferences. Animal Behaviour 29, 628–629 (1981).

41. McNamara, J. M. & Houston, A. I. Risk-sensitive foraging: A review of the theory. Bulletin of Mathematical Biology 54, 355–378 (1992).

42. Caraco, T., Martindale, S. & Whittam, T. S. An empirical demonstration of risk-sensitive foraging preferences. Animal Behaviour 28, 820–830 (1980).

43. De Petrillo, F. et al. Contextual factors modulate risk preferences in adult humans. Behavioural Processes 176, 104137 (2020).

44. Heilbronner, S. & Hayden, B. Contextual Factors Explain Risk-Seeking Preferences in Rhesus Monkeys. Front. Neurosci. 7, (2013).

45. Leblond, M., Fan, D., Brynildsen, J. K. & Yin, H. H. Motivational State and Reward Content Determine Choice Behavior under Risk in Mice. PLOS ONE 6, e25342 (2011).

46. Fehr-Duda, H., Bruhin, A., Epper, T. & Schubert, R. Rationality on the rise: Why relative risk aversion increases with stake size. J Risk Uncertain 40, 147–180 (2010).

47. Webb, J. et al. Foraging Under Uncertainty Follows the Marginal Value Theorem with Bayesian Updating of Environment Representations. 2024.03.30.587253 Preprint at 10.1101/2024.03.30.587253 (2024).

48. van Gils, J. A., Schenk, I. W., Bos, O. & Piersma, T. Incompletely Informed Shorebirds That Face a Digestive Constraint Maximize Net Energy Gain When Exploiting Patches. The American Naturalist 161, 777–793 (2003).

49. Yu, Z. et al. Beyond t test and ANOVA: applications of mixed-effects models for more rigorous statistical analysis in neuroscience research. Neuron 110, 21–35 (2022).

